# Homeostatic Binary Networks: A simple framework for learning with overlapping patterns

**DOI:** 10.1101/2025.10.03.680195

**Authors:** Albert Albesa-González, Claudia Clopath

## Abstract

Memories are rarely stored in isolation: experiences overlap in time and context, leading to neuronal activity patterns that share elements across episodes. While such overlap supports generalization and abstraction, it also increases interference and threatens representational stability. Here we introduce Homeostatic Binary Networks (HBNs), a minimal recurrent framework that combines binary activity, adjustable inhibition, Hebbian learning, and homeostatic plasticity to address these challenges. First, we formalize an Episode Generation Protocol (EGP) that creates compositional episodes with controllable overlap and noise, and define a corresponding semantic structure as conditional probabilities between concepts. We then show analytically and through simulations that recurrent synapses converge to conditional firing probabilities, thereby encoding asymmetric semantic relationships across concepts. These recurrent dynamics enable reliable recall and replay of overlapping episodes without representational collapse. Finally, by incorporating feed-forward plasticity with a neuronal maturity mechanism, output neurons form selective receptive fields in a one-shot manner and refine them through replay, yielding robust unsupervised classification of overlapping episodes. Together, our results demonstrate how simple principles such as neural and synaptic competition can support the stable representation and organization of overlapping memories, providing a mechanistic bridge between episodic and semantic structure in memory systems.

## Introduction

A central goal of memory research is to explain how the brain represents and organizes experiences that are conceptually overlapping and compositional (O’Reilly et al., 2012; Kurth-Nelson et al., 2023). Since the earliest formulations of the hippocampus as a cognitive map (O’Keefe & Dostrovsky, 1971; Moser et al., 2008), to later findings of hippocampal and parahippocampal regions encoding latent environmental variables (Quiroga et al., 2005; Aronov et al., 2017), it has been clear that memories are not stored in isolation. Experiences share elements across time and context, producing neuronal activity patterns that overlap. The meaning of this overlap has been debated: concept-selective neurons in medial temporal lobe (“concept cells”) (Kreiman et al., 2000; Quiroga et al., 2005; Quiroga, 2012; Quian Quiroga, 2023; Rey et al., 2025) suggest that overlap encodes semantic relatedness, while theories of distributed representations argue that overlap reflects conjunctive coding of multiple concepts within episodes (Rumelhart & Zipser, 1985; O’Reilly & Rudy, 2001; Singh et al., 2022; Chrysanthidis et al., 2022).

These overlapping codes pose both an opportunity and a challenge. On one hand, they allow generalization, semantic abstraction, and flexible recombination of experiences (Rogers & McClelland, 2004; Lavenex & Amaral, 2000; Sun et al., 2023). On the other hand, networks learning from correlated patterns suffer from interference, representational collapse, and reduced capacity (Treves & Rolls, 1991; Rolls, 2013; Fung & Fukai, 2023; Halvagal & Zenke, 2023). Biological systems must therefore balance pattern separation with pattern completion (Squire, 2004; Rolls, 2013), enabling distinct episodes to remain accessible while their shared structure accumulates across time.

Another challenge in learning from overlapping patterns comes from a key limitation in most commonly used Hebbian methods. Standard methods such as Hopfield (Hopfield, 1982) and Sparse Hopfield (or Hopfield-Tsodyks) (Tsodyks & Feigel’man, 1988; Gastaldi et al., 2021) Networks, or Boltzmann Machines (Ackley et al., 1985) all rely on symmetric connections. This makes it difficult to represent asymmetric semantic relationships, which are ubiquitous across concepts (for example a *pasta* is tightly related to the concept of *food*, much more than *food* is to *pasta*, in the sense that the second certainly evokes the thought of the first, but the reverse association is much weaker). This highlights the need to expand our current approaches to correlation-based learning to account for how such asymmetric representations might be encoded in the brain.

In this work we introduce *Homeostatic Binary Networks* (HBNs), a minimal recurrent model that addresses these challenges. HBNs combine binary activity, Hebbian learning, and homeostatic plasticity to capture how overlapping patterns can be represented, recalled, and reorganized. Our contributions are fourfold.

1. **Formalizing overlapping input** We propose an *Episode Generation Protocol* (EGP) that generates episodes compositionally, with controllable conceptual overlap and noise. Our episodic input formalizes the hypothesis that overlaps are due to conjunctive coding of multiple concepts across episodes, rather than indicating semantic relatedness. Together with our EGP, we define the *semantic structure* across concepts as a matrix of conditional probabilities between semantic units.
2. **Recurrent weight dynamics** We show that recurrent synapses converge to conditional firing probabilities, therefore connecting them to our definition of semantic structure.
3. **Recall of overlapping patterns without collapse** Using these recurrent dynamics, the network recalls de-noised episodes from partial cues and replays them from noise.
4. **Classification of overlapping patterns with replay and a feed-forward maturity mechanism** Adding plastic feed-forward synapses with a maturity mechanism enables output neurons to form receptive fields in a one-shot manner and refine them gradually, leading to the unsupervised recovery of prototypes and correct classification of overlapping episodes.

## Results

### Episodic generation of overlapping input patterns with semantic structure

Neuronal activity patterns often overlap, and these overlaps have been viewed as signatures of semantic relationship. One interpretation ((Gastaldi et al., 2021; Gastaldi & Gerstner, 2024)) is that overlap reflects *semantic relatedness* between patterns representing different concepts (e.g., *spaghetti* and *fish*; Fig. 1A1). Under this perspective, if two concepts are semantically related their neuronal overlap will be higher. However, this interpretation assumes that each activity pattern encodes only a single concept. Furthermore, this view cannot account for *asymmetric* semantic relationships -for example, *spaghetti* is always food, but food is not always *spaghetti*, yet the overlap between the two is identical in both directions.

**Figure 1:**
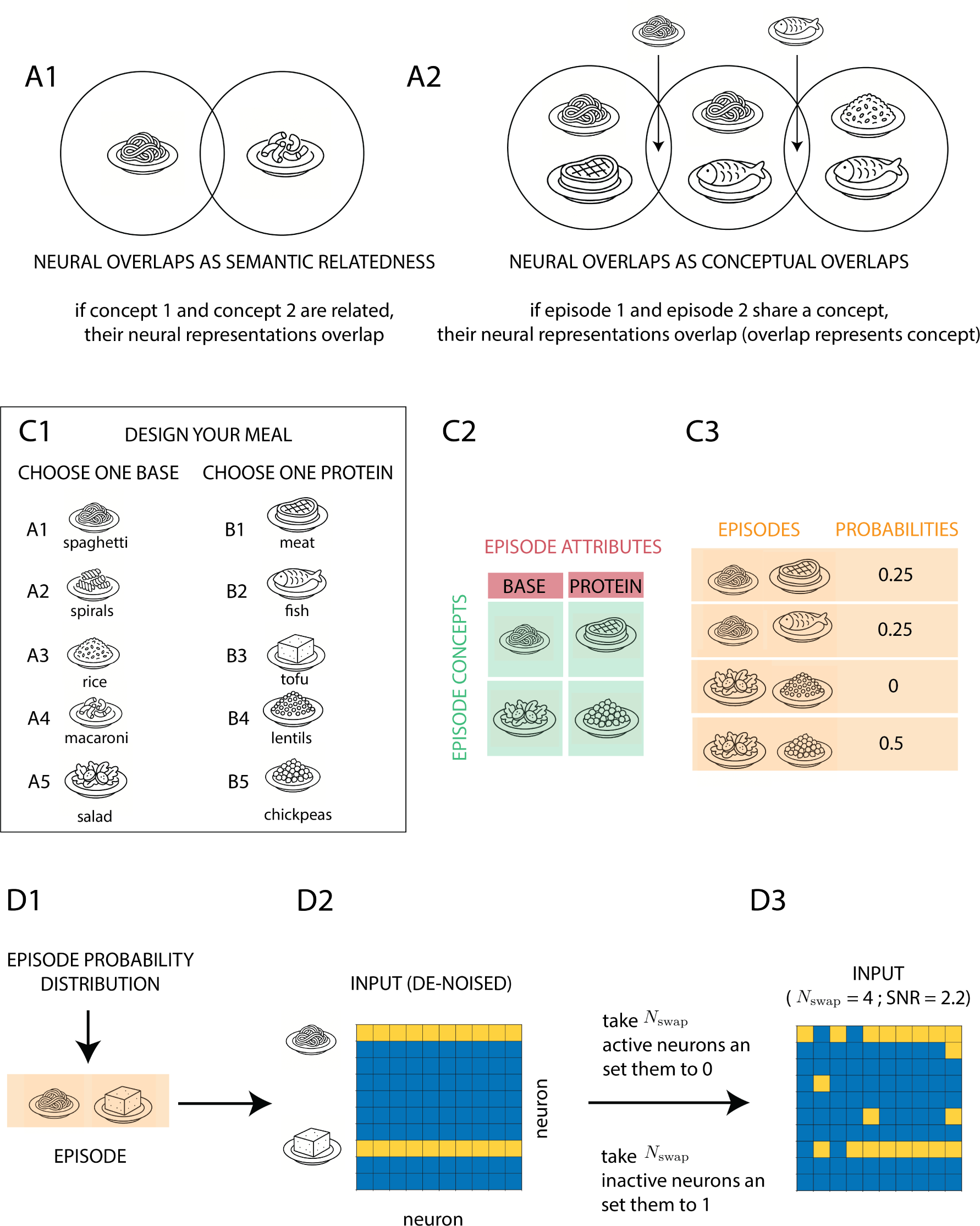
Episodic generation of overlapping input patterns with semantic structure. **A1:** Traditional interpretation of overlap as semantic relatedness, where conceptually similar patterns (e.g., *spaghetti* and *macaroni*) share neurons. **A2:** Alternative interpretation adopted here, where overlap arises from episodes sharing multiple concepts simultaneously. **C1–C2:** Episode Generation Protocol (EGP): episodes are built compositionally by selecting one concept from each attribute (e.g., a base and a protein to form a meal). **C3, D1:** A probability distribution specifies the frequency of each episode, defining the generative structure. **D2:** Each chosen concept activates its associated set of neurons, producing a compositional activity pattern. **D3:** Noise is introduced by swapping the activity of *N*_swap_ pairs of active and inactive neurons.

We instead adopt the interpretation that activity patterns represent full episodes (conjunctions of concepts), and that overlap arises when *episodes share concepts*. Following this, two patterns overlap when a concept is encoded in both (Fig. 1A2), with the overlap corresponding to the representation of the shared concept. This view has been made explicitly (Chrysanthidis et al., 2022) and implicitly (Singh et al., 2022) in previous work. We extend this view by proposing that semantic relatedness might be encoded in the synaptic connections between assemblies of neurons coding for different concepts, which would allow representing asymmetric semantic relationships. Below, we will propose synaptic connections are a natural candidate to represent semantic relationships given that, unlike neuronal overlap, synaptic strength can be asymmetric.

To formalize this view, we introduce an *Episode Generation Protocol* (EGP, Fig. 1C–D). In our EGP, episodes are defined compositionally by selecting one concept from each attribute (e.g., a *base* and a *protein* to form a *meal*; Fig. 1C1–C2). A probability distribution specifies the frequency of each possible episode (Fig. 1C3, D1). Then, every chosen concept activates its corresponding set of neurons, yielding a compositional activity pattern (Fig. 1D2). Noise is introduced by flipping (from 1 to 0) *N*_swap_ active neurons, as well as *N*_swap_ inactive neurons (from 0 to 1, Fig. 1D3, Methods). Defining Signal-to-Noise Ratio (SNR) under this noise implementation is relatively straightforward without relying on continuous variables, maintaining a simplified binary formulation of input (Methods).

This protocol allows naturally defining a *semantic structure* based on the conditional probability of one concept given another (Methods):

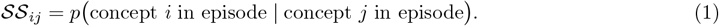

Importantly, this definition can capture asymmetric relationships. For example, let’s imagine that someone always gets *meat* as a protein when *spaghetti* as a base (Fig. 2A). For any other base, the protein is chosen at random. In that scenario, the memory trace evoked by *spaghetti* will be strongly referencing *meat*, whereas the opposite relationship is weaker, since meat can also occur with other bases (Figs. 2B and 2C).

**Figure 2:**
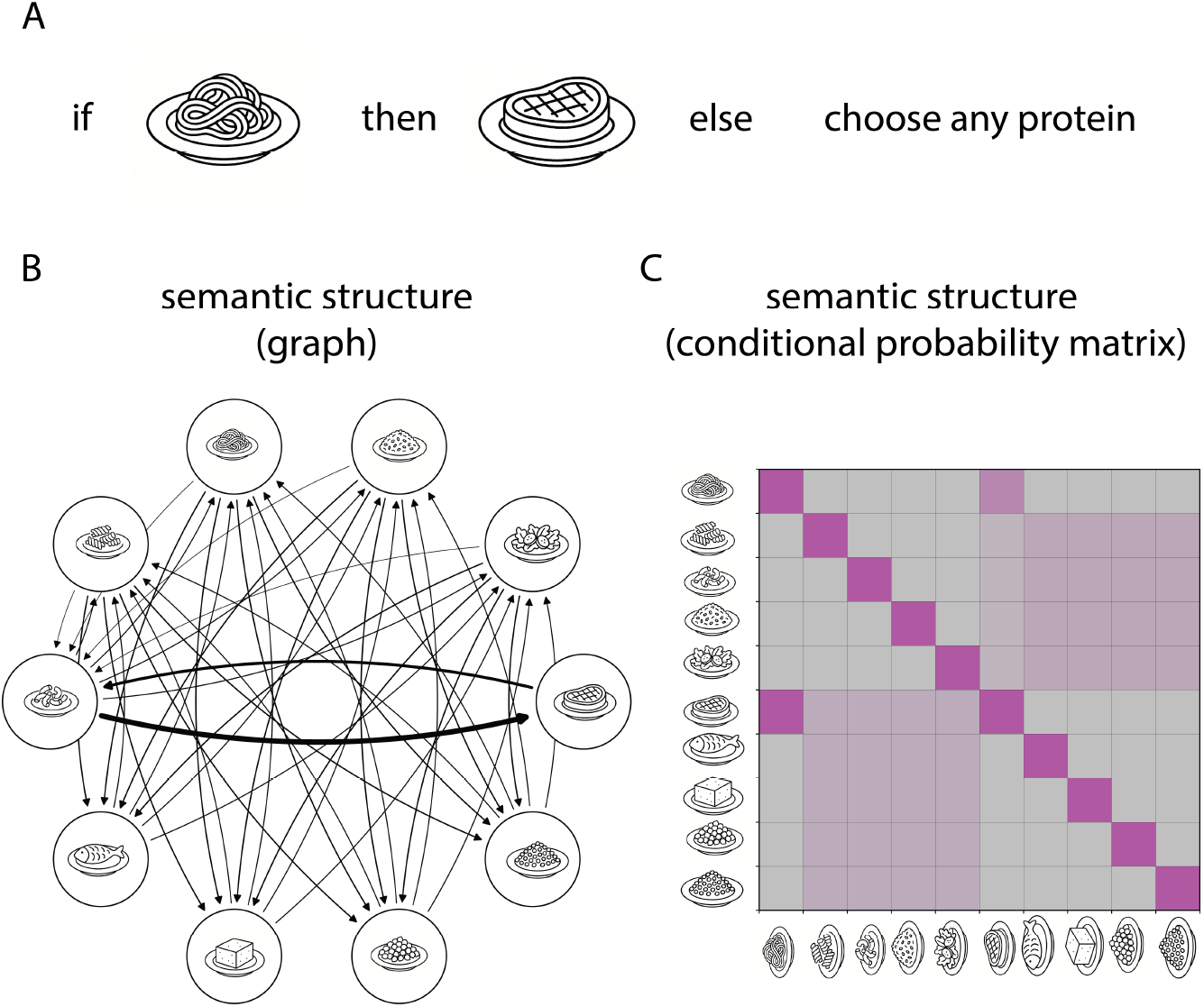
Semantic structure defined by the Episode Generation Protocol (EGP). **A:** Example of asymmetric semantic relationship: *spaghetti* always pairs with *meat*, but *meat* can also occur with other bases, producing stronger association in one direction than the other. **B:** Semantic structure as a directed SS graph with weights representing conditional probabilities of episode co-occurrence. Note the asymmetric relationship between *spaghetti* and *meat*. **C:** Same but in matrix form, with _*ij*_ representing the conditional probability of concept *i* in an episode given concept *j* is in an episode.

### *K-winners-take-all* mechanism generates mild selectivity to highly overlapping patterns

Having formalized a way of generating overlapping patterns with semantic structure, we propose a simple model designed to extract semantics while simultaneously distinguishing episodes that are conceptually overlapping. Our model is a binary neural network equipped with homeostatic mechanisms in both neural and synaptic dynamics, which we call *Homeostatic Binary Network*. The model comprises a hidden recurrent layer *W* ^hidden←hidden^ that maps input to an output layer via feed-forward synapses *W* ^output←hidden^ (Fig. 3A).

**Figure 3:**
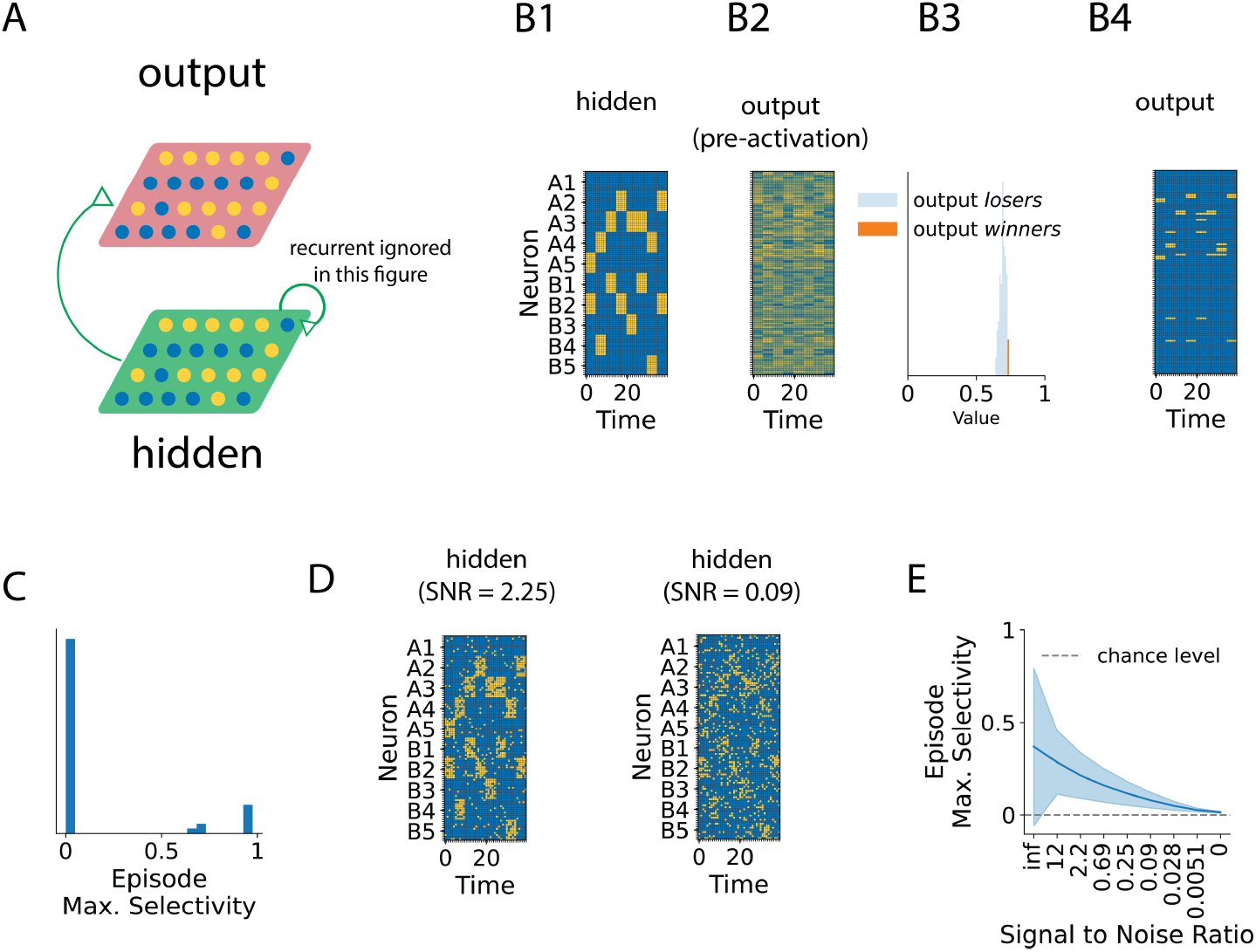
*K-winners-take-all* mechanism generates mild selectivity to highly overlapping patterns. **A:** Model architecture: a hidden recurrent layer projects to an output layer via random feed-forward synapses (recurrent synapses ignored here). **B1:** Example hidden activity generated using the Episode Generation Protocol. **B2:** Pre-activation in the output layer obtained via random feed-forward projection. **B3:** Application of the K-winners-take-all (top-*K*) mechanism, which selects the *K* most active output neurons (orange denotes active neurons (*winners*) and blue inactive neurons (*losers*). **B4:** Post-activation activity of output layer across time, with neurons ordered by episode selectivity. **C:** This competitive mechanism enhances neuronal selectivity to overlapping episodes, even with random connectivity. **D:** Example hidden activity with different Signal-to-Noise Ratio (SNR). **E:** Change in episode selectivity of output neurons with different values of SNR.

We first study the dynamics of the model in the absence of plasticity. We focus on the projection of hidden activity into an output layer via feed-forward connectivity, which is sampled from Gaussian noise (see Methods). To do so, we generate activity in the hidden layer following an Episode Generation Protocol as described in Fig. 1. In particular, we choose input to have 5 possible concepts associated with *A* (*A*_*i*_ ∈ *A*_1_, …, *A*_5_) and 5 with *B* (*B*_*i*_ ∈ *B*_1_, …, *B*_5_). Each *A*_*i*_ (and *B*_*j*_) can be seen as a base (and protein) in the previous example. Episodic statistics are chosen such that pairs of the form (*A*_*i*_, *B*_*j*=*i*_) are sampled half of the time, and the rest -(*A*_*i*_, *B*_*j*≠*i*_)-are equally distributed in the remaining of timesteps. These statistics introduce an imbalance across episodic patterns, making it harder to develop selective neurons across all episodes. Then, input is copied into the hidden layer (Fig. 3B1), which for now ignores recurrent connections. Then, activity in the hidden layer is linearly projected (Eq. 2) to the output layer, resulting in a postsynaptic pre-activation pattern 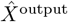 (Fig. 3B2).

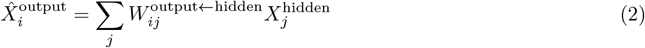

We choose a *K-winners-take-all* (or top-*K*, Eq. (3)) mechanism as mapping from pre-activation to activation values:

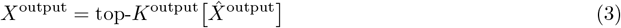

This mechanism takes the pre-activation distribution 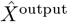 and selects the *K*^output^ neurons with more synaptic input. These neurons set their activity to 1 and the rest are set to 0 (Fig. 3B3 and Fig. 3B4). This activation function can be interpreted as a Heaviside activation with a dynamic threshold, with a biological inspiration in adjustable inhibition (Wehr & Zador, 2003; Vogels et al., 2011). This is one of the most important components of our Homeostatic Binary Network, as it guarantees that the network remains *homeostatic* in its *activity*, by imposing a stable number of active neurons (fixed sparsity) and *binary*. This mechanism maintains the sparsity *a*^region^ of a region

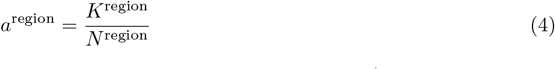

fixed, with *K*^region^ the number of active neurons in a region at a given time, and *N* ^region^ the total number of neurons in the given region. Second, it enforces an amplification of differences in synaptic input, yielding a relatively high neuronal selectivity, to episodes (Fig. 3C), even with non-sparse random connectivity. However, in the presence of noise (Fig. 3D), selectivity is rapidly erased for lower signal-to-noise ratios (Fig. 3E).

These results show that, while a K-winners-take-all dynamics can separate highly-overlapping input patterns, this separation is only partial and not robust to noise. In the following sections, we will explore how these episodes can be better classified using learning in recurrent and feed-forward synapses.

### Recurrent connections converge to conditional firing probabilities, reflecting semantic structure

An input structure like the one presented here contains two sources of overlap across different episodes. On the one hand, (i) episodes are conceptually overlapp4ing, meaning that even in the absence of noise input patterns can contain a high proportion of active neurons (here 50%) if both episodes share one (and only one) concept -such as (*A*_1_, *B*_1_) and (*A*_1_, *B*_2_). On the other hand, (ii) noise creates further spurious correlations across patterns, even when these are conceptually orthogonal (for example (*A*_1_, *B*_1_) and (*A*_2_, *B*_2_))). In this section, we explore how the recurrent network can filter statistical regularities in activity (i), while discarding correlations caused by noise (ii).

To address this, we analyzed the dynamics of recurrent weights under our learning rule. Our learning rule contains two distinct components: one Hebbian

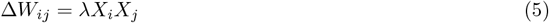

and one homeostatic. Homeostatic plasticity guarantees that either the maximum amount of incoming (∑_*j*_ *W*_*ij*_) or outgoing (∑_*j*_ *W*_*ij*_) connections is capped, which is done via multiplicative normalization (Eqs. (28) and (27) in Methods). Depending on whether the outgoing or incoming sum is normalized, we call it *outgoing homeostasis* or *incoming homeostasis* (respectively).

To get an intuition of how these learning dynamics behave, we turn to analytical methods and solve the learning trajectories of passive (or *non-physical*; neural activity is not affected by synapses) connections with an imposed firing statistical structure, and obtain the following expressions (see Methods):

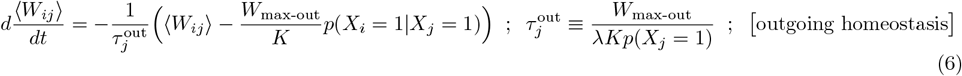

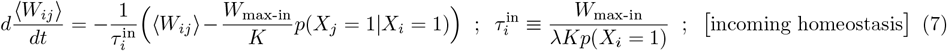

where we have dropped all region superindices (region in hidden, and connectivity is hidden ← hidden), *λ* is the hidden ← hidden learning rate, *W*_max-in/out_ the maximum incoming/outgoing connections, *K* the number of active neurons in hidden, *p*(*X*_*i*_ = 1) the probability of hidden neuron *i* being active and *p*(*X*_*i*_ = 1 |*X*_*j*_ = 1) the probability of neuron *i* being active given neuron *j* is active.

We find that both under outgoing and incoming homeostasis individual weight trajectories converge to a fixed point that depends on conditional firing probabilities. Furthermore, this convergence follows exponential trajectories from initial connectivity, with a time constant (*τ*_*w*_ in Eqs. 6 and 7) that depends on three quantities relevant for biology: (i) the maximum (out/in) connectivity 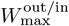, (ii) the learning rate *λ*, and (iii) the pre/post firing rate 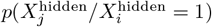. Furthermore, this result connects learning trajectories with the *semantic structure* defined in Fig. 2 (in the case of outgoing homeostasis, assuming each neuron perfectly represents a concept), as well as its transpose (in the case of incoming homeostasis) (Fig. 4B). Another result that follows from this is that, in the presence of asymmetric semantic relationships, weights also become asymmetric. This asymmetry is common across conceptual hierarchies; in our example *tofu* is more semantically related to *protein* than *protein* is to *tofu*.

**Figure 4:**
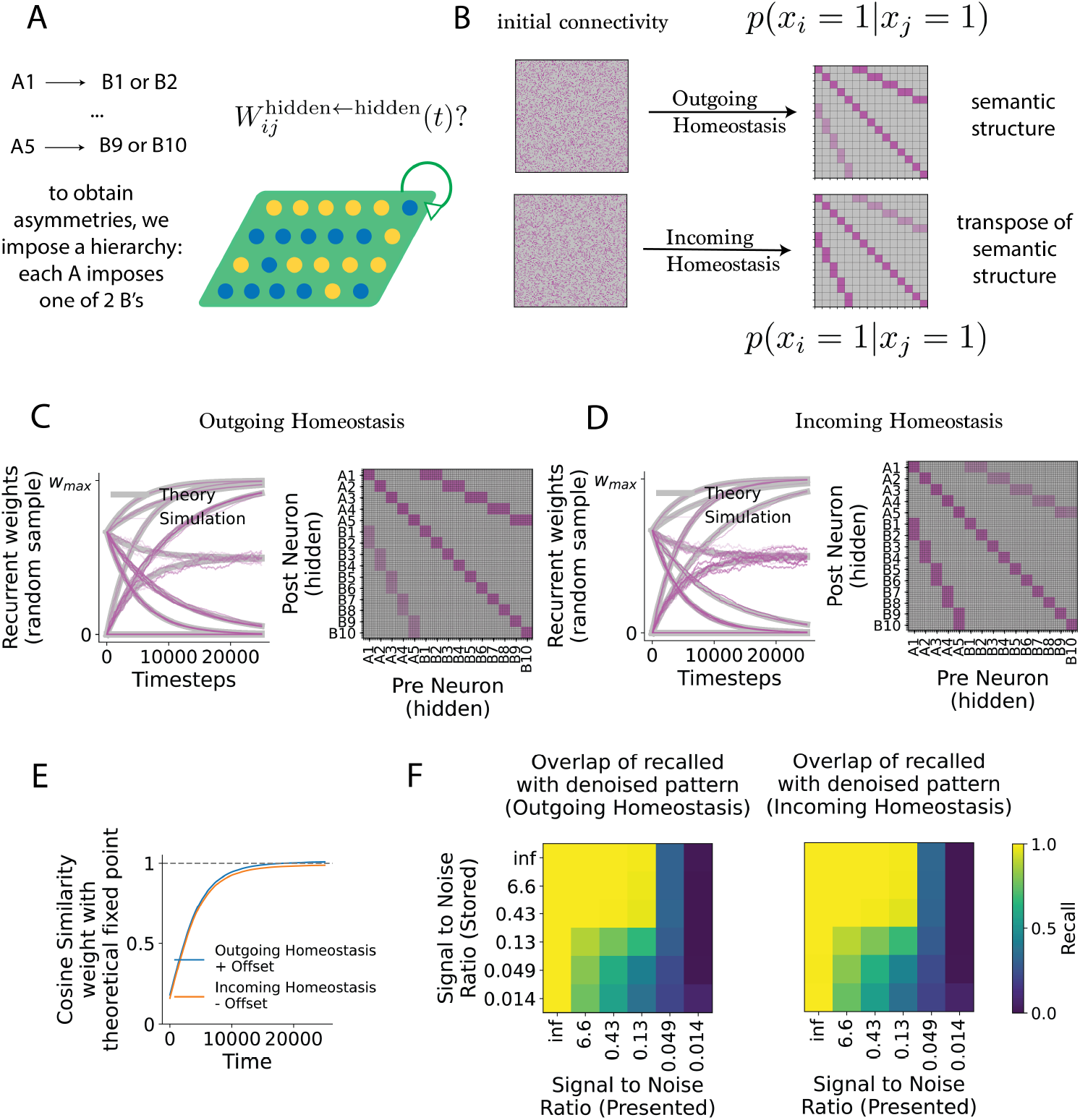
Recurrent connections learn conditional firing probabilities, reflecting semantic structure. **A:** Example Episode Generation Protocol with asymmetric structure, where each base (*A*_*i*_) determines a specific pair of proteins (*B*_2*i*_ or *B*_2*i*+1_). **B:** Analytical prediction of synaptic trajectories under outgoing (left) and incoming (right) homeostasis, showing *W*_*ij*_ converges toward conditional probabilities *p*(concept *i*| concept *j*) and *p*(concept *j*| concept *i*) (respectively). **C–D:** Simulated synaptic dynamics (purple) closely match analytical solutions (gray), with block-like structure emerging in the recurrent weight matrix after learning. **E:** Quantification weight matrix convergence to the theoretical fixed points. **F:** Recall performance: networks trained with noisy overlapping episodes recover de-noised versions, generalizing across different signal-to-noise ratios for both outgoing and incoming homeostasis.

In order to test the property of weight asymmetry in the presence of asymmetric conditional probabilities, we define a new Episode Generation Protocol. In this new protocol, each *A*_*i*_ determines *B*_*j*_ to be either *B*_2*i*_ or *B*_2*i*+1_ (Fig. 4A). This results in an asymmetric semantic structure, and therefore predicts that synaptic connections converge to different fixed points in outgoing and incoming homeostasis (Fig. 4B).

Given that Eqs. (6) and (7) require the condition that synapses are already found at the maximum of their outgoing and incoming connections (respectively), we initialize connections randomly but imposing this condition (Methods). Then, we allow the network to evolve for 25000 timesteps, which corresponds to 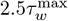 (where 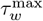 is the slowest time constant across synapses) (Figs. 4C and 4D). A random subsample of synaptic dynamics show a close match between derived (gray) and simulated (purple) trajectories. Furthermore, networks under both outgoing and incoming homeostasis converge (respectively) to the semantic structure and its transpose (compare right panels in Fig. 4B with right panels in Figs. 4C and D), quantified in Fig. 4E).

Next, we hypothesize that the block-like structure obtained in the weight matrix, together with competitive neuronal dynamics during pattern completion, allows recovering distinct episodes, even when these are highly overlapping. Pattern completion is defined as an iterative update of activity in which activity is mapped from hidden to hidden via recurrent connectivity, see Methods). Moreover, as introduced earlier in this section, we test the ability of the network to discard noise-induced correlations, generating de-noised versions of episodes generated with a lower signal to noise ratio (see Fig. 3D). To do this, we trained networks with episodes generated with different signal to noise ratios (SNR (Stored) in Fig. 4F) and then tested recall by presenting new episodes with a different noise level (SNR (Presented) in Fig. 4F). We found that the network can learn the underlying statistical structure in episode generation, while recovering distinct de-noised versions of input even when trained with highly noisy overlapping patterns, and generalizes to highly noisy patterns (Fig. 4F) both for outgoing and incoming homeostasis.

To summarize, we have obtained a closed-form solution for synaptic dynamics in Homeostatic Binary Networks trained with Hebbian and homeostatic plasticity. Our theoretical results, which match trajectories obtained in simulations, show that in our model weights low-pass filter conditional firing probabilities, with a time constant that depends on physiologically relevant parameters. This result connects learning trajectories with the semantic structure of concepts represented by neurons, when semantic structure is defined based on conditional presence of concepts across episodes. Finally, we have demonstrated that the learned connections, in combination with our K-winners-take-all dynamics, allow pattern completion dynamics to retrieve error-corrected episodes, even when de-noised episodes share a high number of active neurons.

### Replay of episodic activity makes neurons in the output layer highly selective to episodes

We have seen how recurrent plasticity enables denoising of episodes during recall, but the network still lacks a stable mapping from hidden activity to distinct episode units. We next hypothesized that we could leverage feed-forward plasticity and episode denoising in order to obtain a high episode selectivity in output neurons. To do this, we introduce a mechanism that combines replay with a learning rate that depends on the *maturity* of output neurons (Fig. 5). We initialize all output neurons to be *immature*, and assign a higher learning rate and excitability to immature units. When immature neurons form an initial rapid receptive field, they become mature. Our assumption is that, just like recurrent connections de-noise corrupted input, replayed activity starting from random patterns would also converge to de-noised episodes (Fig. 5A1). This would mitigate one of the sources of overlap between episodes (noise), simplifying the task of distinguishing between episodes. Then, mildly selective neurons due to random connectivity are activated (Fig. 5A1), and become more selective to the replayed pattern because of their higher learning rate (Fig. 5A2). As replay continues, if the same episode is replayed slower plasticity refines the initially (one-shot) formed receptive field (Fig. 5A3). If a different episode is replayed, a new set of output neurons is activated, following the same process we just described (Figs. 5A2 and 5A3).

**Figure 5:**
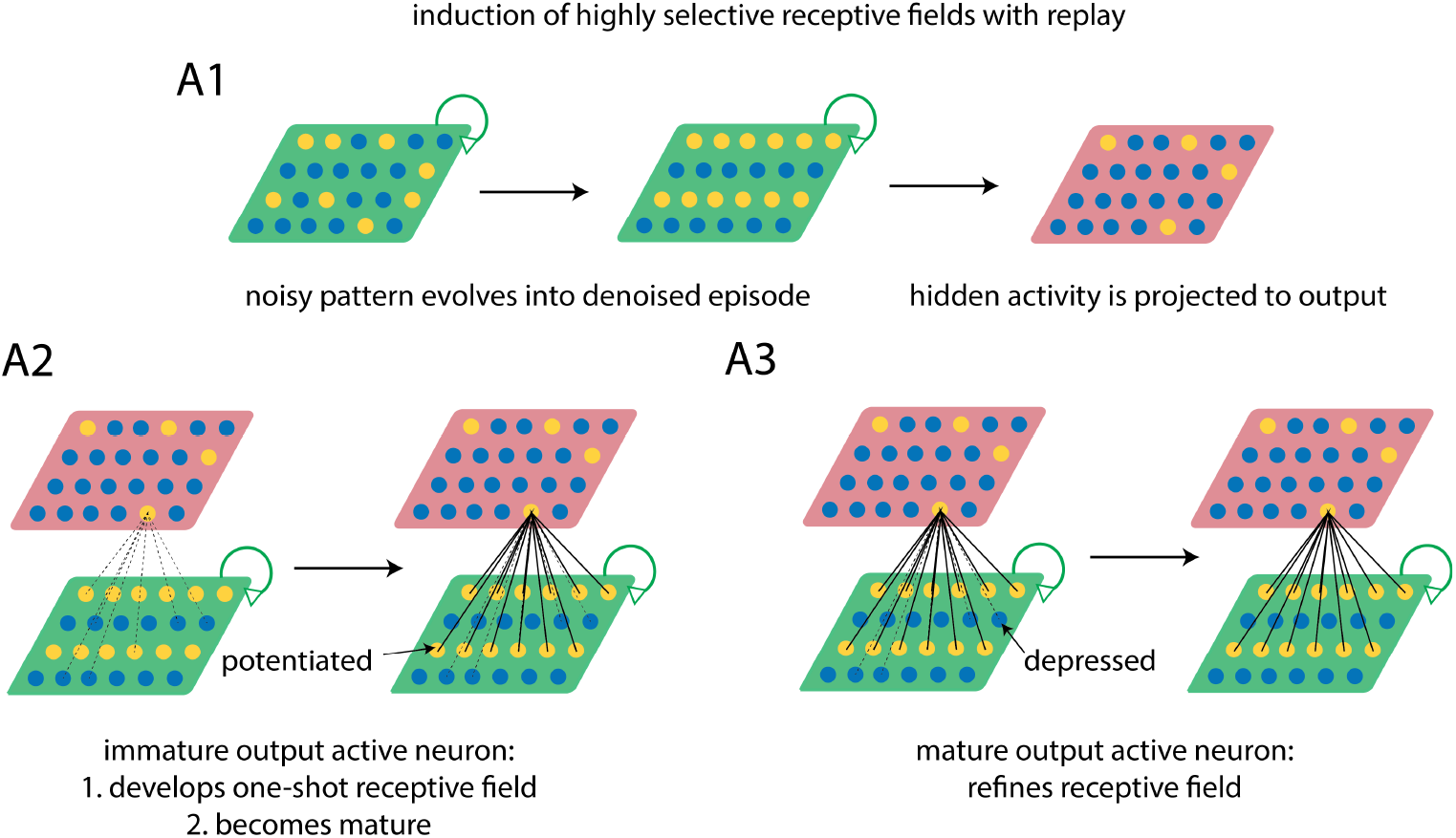
Diagram showing how replay and feed-forward maturation enable formation of episode-selective receptive fields. **A1:** During replay, random patterns converge to de-noised episodes in the hidden layer, which are then projected to the output layer. **A2:** Immature output neurons, with higher excitability and learning rate, become selectively activated by replayed episodes and rapidly form one-shot receptive fields. **A3:** As replay continues, repeated presentations of the same episode refine receptive fields through slower plasticity, while new episodes recruit a different subset of immature neurons.

To test this, we next examined these dynamics in a full network simulation (Fig. 6). At the beginning of training, all output neurons are immature (see output input distribution in Fig. 6A). Pattern completion of noise in the hidden layer recovers a de-noised episode (hidden activity in Fig. 6A), activating a set of output neurons via initially random feed-forward connectivity (see input distribution and activated output in Fig. 6A). These immature neurons form an initial one-shot receptive field of the recovered episode (presynaptic hidden pattern, Fig. 6A, right), because of their higher learning rate. As replay continues, even when an overlapping hidden pattern is recovered (compare hidden activity in Fig. 6A and 6B) a new set of output neurons is activated. Because the initially formed receptive field in Fig. 6A strengthened the connections from *A*_5_ hidden neurons, when pattern (*A*_5_, *B*_5_) is recovered (Fig. 6B), the same output neurons as in Fig. 6A) should in theory be activated again, as the rest of neurons have non-specific random connectivity. This would result in eventually merging the incoming connections from any pattern of the from (*A*_5_, *B*_*j*_), and an eventual representational collapse (which is a known problem with overlapping patterns). However, because neurons active in Fig. 6A became mature and lost their extra excitability, a different set of immature neurons with random connections win the competition (see distribution of pre-activation input of mature neurons selective to overlapping patterns, 6B).

**Figure 6:**
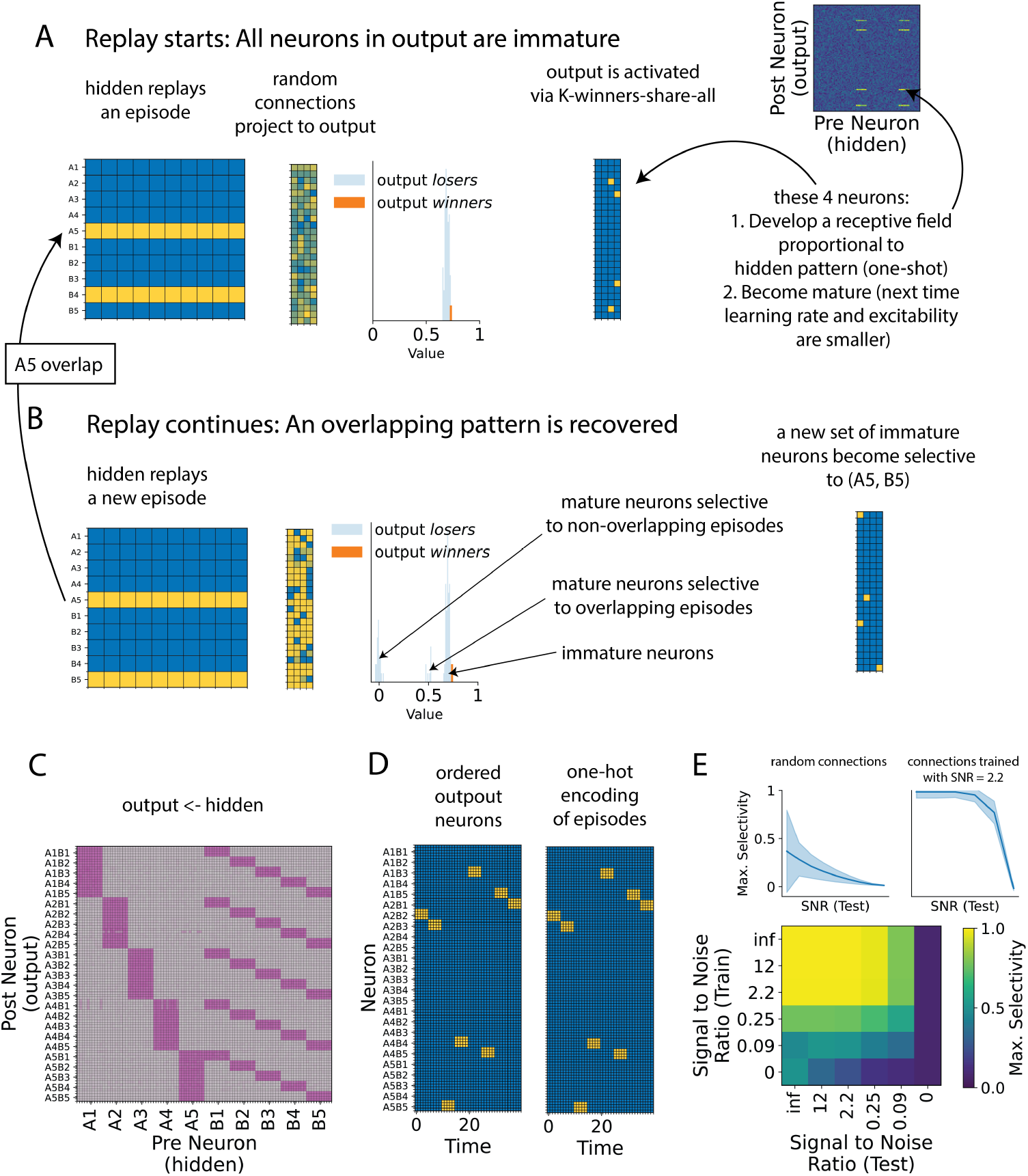
Network simulation shows emergence of episode selectivity and robustness to noise. **A:** Early in training, all output neurons are immature. Replay-driven pattern completion in the hidden layer recovers a de-noised episode, activating a subset of immature output neurons that rapidly form an initial one-shot receptive field. **B:** When an overlapping episode is replayed, a different set of immature neurons is recruited. Previously activated neurons lose excitability as they mature, so they can only compensate the loss of excitability if the same pattern is retrieved (even a pattern that shares half of active neurons activate a completely new set of output neurons. **C:** Over time, output neurons develop stable receptive fields selective to individual episodes. When ordering neurons by their episode selectivity, connectivity matrix shows how output neuron receptive fields (rows) reflect the different de-noised episodes that can be presented to the network. **D:** During wakefulness, noisy inputs are mapped onto the correct episode-selective output neurons, demonstrating robustness of learned representations. **E:** Episode classification accuracy remains high across different noise levels during both training and testing, showing that the network generalizes well even when trained on noisy overlapping input. Top: Comparison of random connectivity (no learning) with learned connections without Signal-to-Noise Ratio (SNR) of 2.2. Bottom: Selectivity across train and test SNRs.

Following this mechanism, at each replay event there are two possibilities: either a previously replayed episode is recovered, inducing slow refinement of connections, or a new episode is replayed, forming a rapid initial receptive field. With time, output neurons develop connections selective to each of the individual episodes (Fig. 6C), which result in neurons firing selectively depending on the presented episode, even during wakefulness (noisy patterns are presented). We test how the network generalizes the learned representations to increasing levels of noise during testing (SNR (Test) in Fig. 6E), for different levels of noise during training (SNR (Train) in Fig. 6E). We show how episodic accuracy (episode classified based on episode selectivity) after training remains high largely independent of noise during training (up to a Signal to Noise Ratio of approximately 0.25) and generalizes well for increasing levels of testing noise (Fig. 6E).

Together, these results show how recurrent replay and feed-forward maturation jointly allow the emergence of episode-selective receptive fields, enabling the network to classify overlapping episodes correctly.

## Discussion

We introduced *Homeostatic Binary Networks* (HBNs) as a minimal framework for learning with overlapping inputs. First, we formalized an *Episode Generation Protocol* (EGP) that produces compositional episodes with controllable overlap and noise, and defined semantic structure as a conditional probability matrix across concepts. Second, we showed analytically and numerically that recurrent synapses converge to conditional firing probabilities, thereby encoding stable co-activation statistics while discarding spurious correlations. Third, we demonstrated that recurrent plasticity enables recall and replay without collapsing overlapping patterns into merged attractors. Finally, we showed that plastic feed-forward synapses with a maturity mechanism allow output neurons to acquire receptive fields and classify overlapping episodes in an unsupervised manner. Together, these results suggest that simple principles but well orchestrated —Hebbian plasticity and homeostatic competition—are sufficient for robust recall, replay, and prototype learning in overlapping environments.

Our input generation and concept learning frameworks can reconcile two apparently distinct views regarding the origin and semantic role of neuronal overlaps in biology. One interpretation of overlap across patterns is that they indicate semantic relatedness (Gastaldi et al., 2021; Gastaldi & Gerstner, 2024). This interpretation assumes neuronal activity patterns represent a single concept, and two patterns have a higher overlap when the concepts are semantically related. A different view taken in previous work (Singh et al., 2022; Chrysanthidis et al., 2022; Kang & Toyoizumi, 2023, 2024), assumes that the overlap itself is precisely what represents the concept. While we have positioned our work in the second interpretation, we propose that the two can be reconciled when comparing concepts that share a third concept towards which they have a very high semantic structure. Our results show that networks can form higher-order concepts as a combination of primitive concepts (for example concept (*A*_1_, *B*_1_) from concept *A*_1_ and concept *B*_1_, Fig. 6). If one looks at the final network representation, concepts (*A*_1_, *B*_1_) and (*A*_1_, *B*_2_) -which are found in the output layer-formally have completely orthogonal representations. However, in practice, concept *A*_1_ will co-activate whenever (*A*_1_, *B*_1_) or (*A*_1_, *B*_2_) are active, inducing an effective persistent neuronal overlap between concept (*A*_1_, *B*_1_) and concept (*A*_1_, *B*_2_) (therefore matching the experimental observation that neural patterns that are semantically related (in this case both are associated to *A*_1_) have a higher overlap. It should be noted, however, that in our model the strict representations of (*A*_1_, *B*_1_) and (*A*_1_, *B*_2_) do not overlap, and the effective overlap is a result of their semantic structure. This hypothesis could be tested, for example, by measuring overlaps across multiple semantically-related concepts (for example, is the overlap across different concepts related to *food* consistently similar?)

Both our Episode Generation Protocol (EGP) and our network model are directly related to Potts model of semantic representations (Kropff & Treves, 2005). The *features* of our EGP can be seen as the different local modules present in Potts models, and the different *concepts* associated to each attribute as the different values each Potts local module can be found in. Similarly, Homeostatic Binary Networks can be seen as a mechanistic implementation of the Potts model. Our *K-winners-take-all* dynamics, together with the learned block-like recurrent connectivity, effectively enforce each network subregion activity to settle in one of the different blocks of recurrently connected neurons. On the other hand, Homeostatic Binary Networks build upon long-standing ideas like Hopfield networks (Hopfield, 1982) and competitive learning (Rumelhart & Zipser, 1985). Classical attractor networks demonstrated how Hebbian synapses stabilize patterns (Hopfield, 1982), but interference from correlated patterns limits their capacity (Treves & Rolls, 1991; Rolls, 2013). Here, we have shown how adjustable inhibition can lead to stable pattern recovery even with patterns with a 50% overlap, which was previously reported to unavoidably lead to representational collapse (Gastaldi et al., 2021).

This model incorporates outgoing homeostasis, which has also recently been included in (Fung & Fukai, 2023). While their work focuses on learning in feed-forward connections, competition for presynaptic resources (outgoing homeostasis) was already found to help learning with overlapping patterns. Another novelty of our model is the combination of replay and neuronal maturity in order to develop receptive fields selective to highly overlapping episodic patterns. Here, replay takes a role as an initial de-noising step, eliminating uncorrelated overlaps that might have been presented during training. This is in contrast with most replay mechanisms, that are instead faithful re-instantiations of awake activity (McClelland et al., 1995; Sun et al., 2023). Even after de-noising, patterns have persistent high overlaps when episodes share one out of two concepts in their representations. We have proposed a neuronal maturity mechanism that allows discrimination of these episodes. Neurogenesis-like maturity mechanisms have been modeled before as changes in plasticity excitability (Gozel & Gerstner, 2021), and has also been observed experimentally (Aimone et al., 2011). Here, this is combined with *K-winners-take-all* dynamics, which enforces a very high degree of specificity in plasticity-inducing neuronal firing. The proposed induction of initial one-shot receptive field is reminiscent of Behavioural Timescale Synaptic Plasticity (BTSP, (Bittner et al., 2017)). While here (concept) place field induction depends on postsynaptic activity and neuronal maturity (in contrast to BTSP, which depends on entorhinal signals (Grienberger & Magee, 2022)), our model highlights the computational advantage of forming initially strong receptive fields in episodic pattern separation.

Homeostatic Binary Networks are deliberately simplified. One of the burdens in modeling neural systems consists in finding the right set of network parameters that stabilize neural activity and learning (e.g., stability is often hard and may require fine-tuning) (Sussillo & Abbott, 2009; Laje & Buonomano, 2013; Abeysuriya et al., 2018; DePasquale et al., 2018). Using *K-winners-take-all* (Maass, 2000) facilitates neural-dynamics stabilization by abstracting the usually implicit process of neural competition and excitatory–inhibitory balance, while directly imposing a desired level of sparsity. The proposed activation mechanism is scale-free, with the results being invariant to synaptic and excitability scaling. This avoids relying on parameter tuning or solving mean-field approximations of the neural dynamics. Similarly, we have shown that the fixed points of the learning dynamics are independent of the learning rate, with this parameter only influencing the timescale in which the fixed point is reached.

However the model presents several limitations, especially related to asymmetric temporal correlations. In this sense, the network acts independently on activity snapshots, and cannot capture sequences of neural activity. Similarly, replay of activity is also limited to replay of static patterns, rather than temporally extended replay of behaviorally-related neuronal sequences. Future work could explore how, for example, representing temporally averaged input could overcome these difficulties, connecting at each timestep the past with the present. Finally, our semantic structure relies on pairwise conditional probabilities, which may not capture higher-order dependencies seen in cortical representations.

In conclusion, Homeostatic Binary Networks provide a minimal yet effective framework for representing and organizing overlapping episodes. By combining Hebbian and homeostatic plasticity with competitive neuronal dynamics, the model captures conditional structure, enables reliable recall without collapse, and supports the emergence of selective receptive fields through replay and maturation. These results demonstrate that simple principles of neural and synaptic competition are sufficient to achieve stable recall, replay, and classification of overlapping patterns, offering a clear mechanistic account of how episodic and semantic structure can be jointly sustained.

## Methods

### Episode Generation Protocol

The Episode Generation Protocol (EGP) is a generative process that samples episodic activity with a pre-defined semantic structure.

#### Episodes, episode attributes and episode concepts

All *episodes* share a structure, such that every episode contains one *episode concept* per *episode attribute*. This means that, given episode attributes *A* and *B*, with each episode attribute representing a set of episode concepts *A*_*i*_ ∈ *A, B*_*j*_ ∈ *B*, then the set of all possible episodes is defined as:

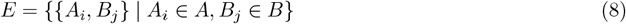

In other words, an episode *e* ∈ *E* is a collection of episode concepts *A*_*i*_, *B*_*j*_ such that element *A*_*i*_ belongs to episode attribute *A* and element *B*_*j*_ belongs to episode attribute *B*. In this study we have fixed *A*_*i*_ to have 5 possible values *A*_*i*_ ∈ {*A*_1_, *A*_2_, *A*_3_, *A*_4_, *A*_5_} and likewise for *B*_*j*_ ∈ {*B*_1_, *B*_2_, *B*_3_, *B*_4_, *B*_5_} (except for Fig. 4, where we have *B*_1_ to *B*_10_).

### Episode Probability Distribution

As a generative process, our EGP samples episodes as a previous step to sampling sensory input. We do this by fixing a probability distribution over episodes:

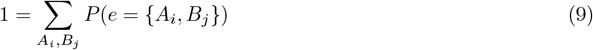

such that *P* (*e* = {*A*_*i*_, *B*_*j*_}) ≡ *P* (*i, j*) imposes the likelihood of an episode with latents (*A*_*i*_, *B*_*j*_) to be sampled when generating episodes.

### Episode-Input Mapping

Ultimately, the EGP samples neural activity. For this reason, one also has to define how a sampled episode is mapped into sensory input *X*^input^. Every concept *c* in attribute *A* ∪ *B* has a set of associated neurons input_c_ in *X*^input^ such that, given an episode *e*

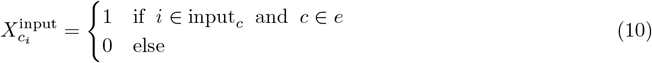

In order to define input_c_, we specify the number of attributes *N*_attributes_, in this case 2 (*A* and *B*). Then, we define the number of concepts per attribute (|*A*| and |*B*|, here 5 and 5 (except in Fig. 4, where is 10), from this follows the fact the fact that *A*_*i*∈[1,5]_ and *B*_*j*∈[1,5]_ —[1, 10] in Fig. 4). Then, we fix the number of neurons per concept 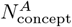 and 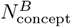, and define input_c_ as:

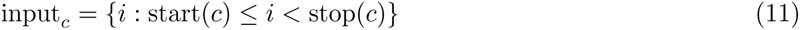

with start 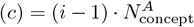 if *c* = *A*_*i*_ ∈ *A* and start 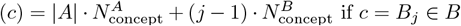, and similarly stop 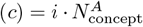 if *c* = *A*_*i*_ ∈ *A* and stop 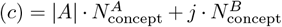 if *c* = *B*_*j*_ ∈ *B* (i.e.we generate a list of one-hot encodings for each concept *c*).

#### Noise

To account for variability between different presentations of the same episode (not all episodes, even though they contain the same concepts, will be exactly the same), we further add some stochasticity by randomly picking *N*_swap_ inactive neurons and *N*_swap_ active neurons, and then flipping their activity (a total of 2*N*_swap_ neurons randomly change their activity). This ensures that activity sparsity in the sensory layer is maintained (the number of flips from 0 to 1 is the same as the number of flips from 1 to 0).

#### Signal-to-Noise Ratio

In order to obtain the Signal-to-Noise Ratio (SNR) of a generated pattern, we first define the signal as the expected overlap (number of common neurons) of the corrupted pattern with the original, and subtract the expected overlap between two randomly generated patterns of *K*^region^ active neurons across *N* ^region^ neurons.

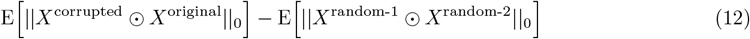

Because we fix that *N*_swaps_ active (in the original pattern) flip from 1 to 0, the first term is always *K*^region^ − *N*_swaps_, while the second corresponds to the expected value of a hypergeometric distribution, given by (*K*^region^)^2^*/N* ^region^. Then, we define the noise to be the number of non-common neurons between the corrupted and the original pattern, which is 2*N*_swaps_, giving:

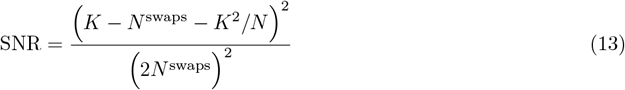

#### Semantic Structure of an EGP

We refer to *semantic field theory* (Bussmann et al., 2006) in order to define what is the *semantic structure* of an EGP. According to this theory, the meaning of a word is not isolated but dependent on its relation to the rest of the words. While our task is not one of language, we can use this same paradigm to define the *meaning* of episode concepts. In this sense, the meaning (semantics) of our episode concepts depends on how they are related to the rest (also note this is how semantics are established in a language dictionary, where each concept is defined via its relationship with the rest of concepts).

Following this proposal, we use the conditional probabilities of being present in an episode between episode concepts as a proxy for the semantic structure of an EGP. In other words, extracting the semantics of an EGP is equivalent to extracting how likely is one episode concept *y* to be present in an episode *e* if an episode concept *x* is also present. We define the Semantic Structure 𝒮𝒮 of an EGP with *N* ^concepts^ different concepts *c*_*i*_, as as a *N* ^concepts^ × *N* ^concepts^ matrix of the form:

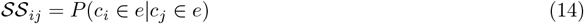

### Circuit Model

#### Architecture and nomenclature

##### Regions and neural activity

The circuit model contains 2 main regions: hidden, and output. In turn, hidden layer is split into two sub-regions: (*A* and *B*). We use *X* to denote the neural activity, which is written as *X*^region^ (for example *X*^hidden^) when the region is to be made explicit. The same follows for pre-activation (synaptic) input, which is denoted by 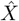 (or 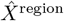). The activity of neuron *i* in a region is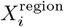. The number of neurons in a region is denoted by *N* ^region^ (e.g. *N* ^hidden^).

##### Connectivity

Connectivity matrices connecting a region pre to a region post take the form *W* ^post←pre^, and a synapse connecting neuron *j* in region pre with neuron *i* in region post is 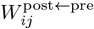. For example, 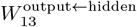 represents the connection from neuron 3 in hidden to neuron 1 in output. Connectivity is classified as either feed-forward (connecting two different regions), which corresponds to *W* ^output←hidden^, or recurrent (connecting a region with itself), which corresponds to *W* ^hidden←hidden^. Given a connectivity matrix *W* ^post←pre^, we call receptive field of a postsynaptic neuron *i* 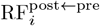 the vector representing the connections from region pre to postsynaptic neuron *i*

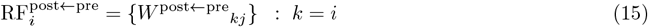

##### Feed-Forward connectivity initialization

Feed-forward connections are initially sampled from a Gaussian distribution with mean 0 and standard deviation 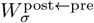 as a network parameter.

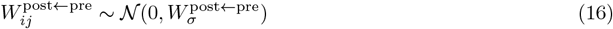

##### Recurrent connectivity initialization

In Fig. 4, *W* ^hidden←hidden^ is initialized randomly with the condition that the sum of the outgoing weights (in the case of outgoing homeostasis) or incoming weights (in the case of incoming homeostasis), matches the maximum allowed by 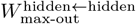 and 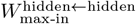 (respectively). We describe how this is done in the case of incoming homeostasis, which can be extended to outgoing homeostasis by taking the transpose of such initialization. First, we think of recurrent connections as feed-forward from hidden to hidden, and then impose in the corresponding receptive fields a specific size:

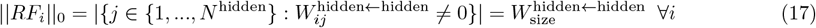

and sum:

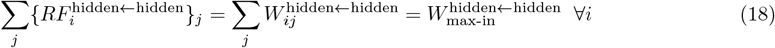

where ||*RF*_*i*_ ||_0_ denotes the *l*_0_ norm (which counts the number of non-zero entries), |*A*| the number of elements in set *A*, 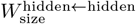 is a network parameter determining the number of non-zero incoming connections at every neuron, and 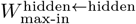 another network parameter determining the maximum sum of incoming connections a postsynaptic neuron has. In order to achieve both conditions, we generate *W* ^hidden←hidden^ by randomly sampling each receptive field *i* independently, following:

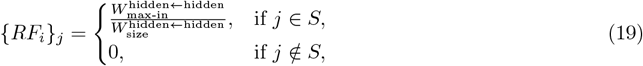

with *S* a random subset of the presynaptic entries:

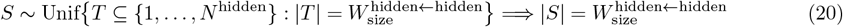

##### Mature and immature neurons

Neurons in the model can be either *mature* or *immature*. The intuition is: do these neurons contain meaningful representations via input received from another region (mature) or are mostly non-selective and random (immature)? More details on the dynamics and plasticity of mature and immature neurons can be seen below, but to summarize: (i) immature neurons have higher excitability during feed-forward input processing and (ii) immature neurons have a higher learning rate in its incoming feed-forward connections. The intuition is that by having higher excitability, during receptive field formation in feed-forward synapses, immature neurons win the competition for postsynaptic activity only if the presynaptic pattern is too far from the receptive fields that have already formed. By doing this, if a presynaptic pattern is similar to previously formed representations, it slightly changes the already formed representations. If not, a set of immature neurons win the competition, develop receptive field due to higher plasticity, and then become mature. This allows one-shot formation of new postsynaptic representations of presynaptic input when an input pattern is too different from those presented to the network before, and then refining these representations with a slower learning rate. Maturity in a region is denoted by IM^region^, where 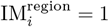 if neuron *i* in the region is immature, and 0 otherwise.

### Network Operations

#### Neuronal activation

Synaptic input is mapped to neural activity via a *K-winners-take-all* mechanism, in which the *K* neurons with the highest pre-activation input 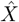 have their activity set to 1, and the rest to 0. This can be written as

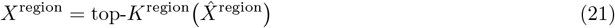

#### Feed-forward input processing

Input processing is computed for a single pre and post region at a time, with pre-activation input 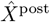 obtained via

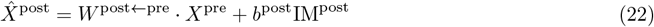

*b*^post^ is the extra excitability that immature neurons have in region post.

#### Pattern completion (recurrent input processing)

Given a recurrent layer, pattern completion is defined as an iterative update on neural activity *X*^region^ that is initiated in an initial state 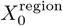:

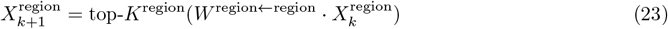

with *W* ^region←region^ the region recurrent connectivity matrix, and *k* a timestep operating on a smaller timescale, than the overall timescale *t*. This means that within a timestep *t*, there are pattern complete iterations iterations on temporal index *k*.

#### Hebbian Learning

The connection between a presynaptic neuron *j* and a postsynaptic neuron *i* subject to Hebbian learning is updated as:

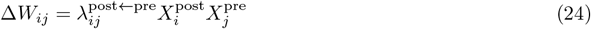

for a presynaptic region pre and a postsynaptic region post, where 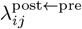 is the learning rate from neuron *j* in region pre to neuron *i* in region post.

#### Maturity-dependent learning rate (feed-forward connections)

In order to combine one-shot and statistical learning in feed-forward synapses, the learning rate of feed-forward (post ≠ pre) connections depends on the *maturity* of the postsynaptic neuron, which is given by IM. Neurons start in an *immature* state IM_*i*_ = 1, allowing them to form relatively selective initial receptive fields. Once this initial receptive field is formed, the learning rate is reduced to capture the statistical regularities of the presynaptic patterns driving the postsynaptic neuron (the neuron is *mature*, IM_*i*_ = 0). To do this, we define a quick learning rate

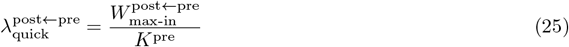

where again 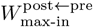 is a network parameter determining the maximum sum of incoming connections the matrix *W* ^post ← pre^ has for every postsynaptic neuron (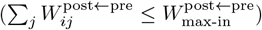). Then, at each Hebbian update, we have

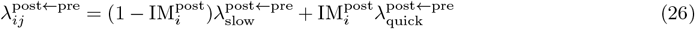

where 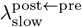 is a network parameter indicating the slow learning rate (used throughout all simulations except for the first receptive field formed at each postsynaptic neuron). Note how for an initial (before maturity) driving presynaptic pattern *X*^pre^ this guaranties that the initial receptive field formed is *X*^pre^ scaled such that the sum of incoming connections is exactly equal to 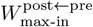, as the *K*-winners-share-all mechanism imposes exactly *K*^pre^ active neurons.

#### Homeostatic Plasticity

We use multiplicative normalization that guarantees, for every weight matrix *W* ^post←pre^, the total sum of incoming or outgoing connections is capped to a certain value. We define *incoming homeostasis* as:

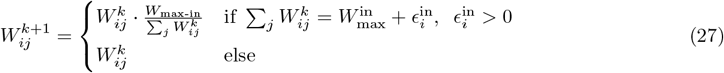

where 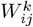 denotes the value of synapse *ij* before applying homeostasis and 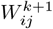 the value after applying homeostasis (for readability, we simply write *W* ≡ *W* ^post←pre^). Similarly, we have *outgoing homeostasis* to be defined by:

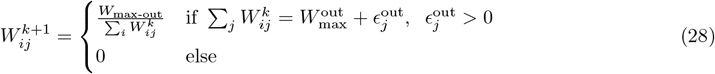

#### Replay

We define Replay as a learning mechanism involving a pre and a post region, such that *W* ^post←pre^ extract statistical structure from *W* ^pre←pre^ connections. To do this, we first pattern complete an initial presynaptic pattern 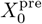, defined as a Gaussian random vector of the same size as the pre region:

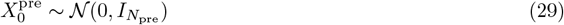

where *N*_pre_ is the size of region pre and 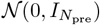 is a multivariate Gaussian distribution with 0 mean and identity covariance. This initial random pattern is pattern completed to obtain *X*^pre^, and then projected to the post region, to give *X*^post^. Then, Hebbian and homeostatic plasticity from post to pre region follow.

**Table 1:**
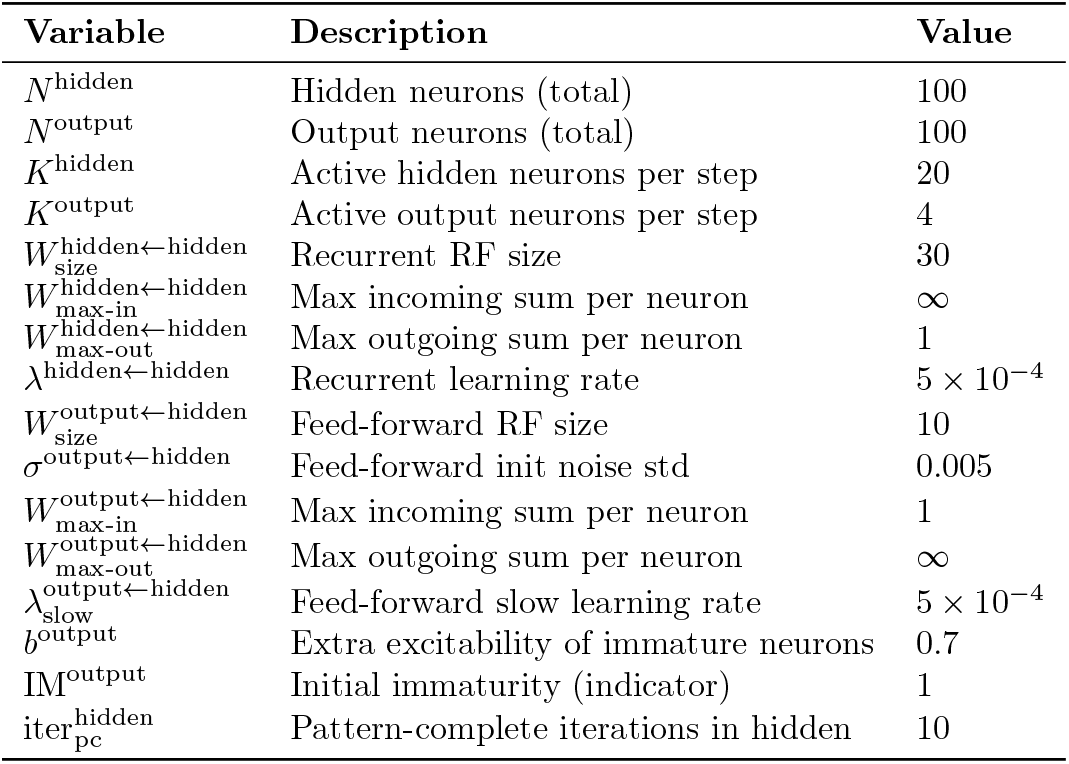
Default model parameters.

**Table 2:** Figure 2 parameters (different than default)

| Variable | Description | Value |
| --- | --- | --- |
| Figs. 2B-2C |  |  |
| $\text{Day}_T$ | Day length | 40 |
| Fig. 2D1 |  |  |
| $N_{\text{swap}}$ | Number of swapped neurons | 4 |
| Fig. 2D2 |  |  |
| $N_{\text{swap}}$ | Number of swapped neurons | 4 |
| Fig. 2E |  |  |
| $N_{\text{swap}} \text{ range}$ | List of simulated number of swaps | 0 to 16 (step = 2) |
| $\text{seed range}$ | List of simulated seeds | 0 to 10 (step = 1) |

**Table 3:** Figure 3 parameters (different than default)

| Variable | Description | Value |
| --- | --- | --- |
| $N^{\text{hidden}}$ | Number of neurons in hidden | 150 |
| $N^{\text{hidden-A}}$ | Number of active neurons in hidden $A$ | 50 |
| $N^{\text{hidden-B}}$ | Number of active neurons in hidden $B$ | 100 |
| $K^{\text{hidden-A}}$ | Number of active neurons in hidden $A$ | 10 |
| $K^{\text{hidden-B}}$ | Number of active neurons in hidden $B$ | 10 |
| $N_B$ | Number of concepts in $B$ | 10 |
| $P(A_i, B_j)$ | Probability distribution | 1/10 if $i = j//2$ else 0 |
| Figs. 3D and 3F2 |  |  |
| $W_{\text{max-out}}^{\text{hidden} \leftarrow \text{hidden}}$ | Max. Outgoing | ∞ |
| $W_{\text{max-in}}^{\text{hidden} \leftarrow \text{hidden}}$ | Max. Outgoing | 1 |
| Figs. 3F1 and 3F2 |  |  |
| $N_{\text{swap}}$ range | List of simulated number of swaps | 0 to 24 (step = 4) |
| seed range | List of simulated seeds | 0 to 10 (step = 1) |

**Table 4:** Figure 4 parameters (different than default)

| Variable | Description | Value |
| --- | --- | --- |
| Fig. 4E |  |  |
| $N_{\text{swap}}$ range | List of simulated number of swaps | 0 to 16 (step = 2) |
| seed range | List of simulated seeds | 0 to 10 (step = 1) |

### Data Analysis

#### Neuronal Selectivity

Given a vector *X* that contains neuronal activity across T timesteps -with dimensions (*T, N* ^neurons^)- and one that contains certain latent variables *Z* -with dimensions (*T, N* ^latents^), we first normalize:

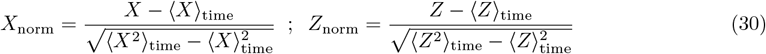

And then obtain the matrix

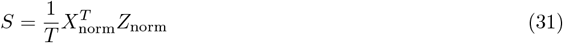

which, for neuron *i* and latent *j* contains their correlation as:

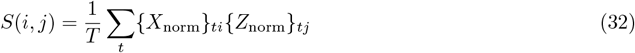

The Maximum Selectivity of neuron *i* is the maximum selectivity value of neuron *i* across all latents *j*:

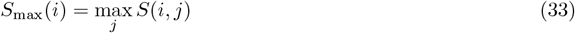

### Recall

In order to evaluate recall, we sample *T* ^train^ episodes from our Episode Generation Protocol, generating *X*^train^ with dimensions (*T* ^train^, *N* ^hidden^). Signal-to-Noise Ratio during training -SNR (Train)-comes from the number of swaps in the generation of this input. Then, we train recurrent connections in hidden with Hebbian and homeostatic plasticity, fixing 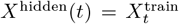. After doing this, we generate *T* ^test^ samples without noise *X*^test-uncorrupted^, and the corresponding noisy version, *X*^test-corrupted^, according to SNR (Test). For each test timestep, we fix 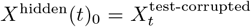, and pattern complete (Eq. (23)) to obtain *X*^hidden^(*t*)_num_iterations_. We define recall as the average cosine similarity of *X*^hidden^(*t*)_num_iterations_ with 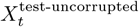.

### Closed-form solutions to learning trajectories

Here, we obtain the learning dynamics of a weight *W*_*ij*_. The intuition is very similar to that presented in Rumelhart and Zipser (1985), which establishes the fixed points of a feed-forward network with incoming homeostasis, extending it to the full weight trajectories, and for both incoming and outgoing homeostasis.

We assume a post and a pre region (both could be the same in the case of a recurrent layer), but drop region indices unless necessary, yielding *W* ≡ *W* ^post←pre^, *λ* ≡ *λ*^post←pre^. We similarly simplify the expressions for neuronal statistics as follows:

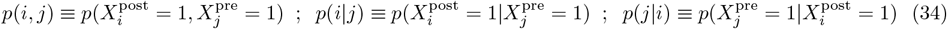

Furthermore, we define the total outgoing (incoming) sum of *W* in neuron *j* (*i*) as at time *t* as:

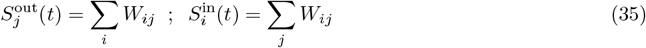

#### Only Hebbian learning

In the absence of homeostasis, in the case that there is outgoing homeostasis and 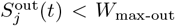, or in the case that there is incoming homeostasis and 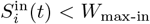, synapse *ij* is only subject to Hebbian learning. The mean trajectory of *W*_*ij*_, can be trivially obtained assuming an initial condition 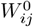:

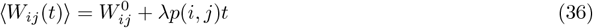

given the only source of plasticity is Hebbian (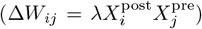) and, on average, this update will be applied in a fraction *p*(*i, j*) of the time.

#### Outgoing homeostasis

Now, let’s consider the case where at *t* = 0, 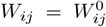, and 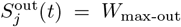, in the presence of outgoing homeostatic plasticity. Given that the only source of potentiation is Hebbian learning, the condition 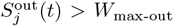 can only be met up upon future firing of neuron *j*. When that happens, according to Eq. (28), we have

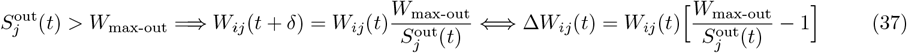

where *W*_*ij*_(*t* + *δ*) indicates the value after homeostasis, which we assume to be instantaneous (in simulations happens within the same simulation timestep).

Eq. (37) can be expanded by rewriting 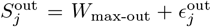 (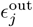 is guaranteed to be positive because 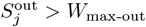) and then using 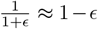 for small *ϵ*, the change due to homeostatic plasticity in synapses outgoing from neuron *j* is:

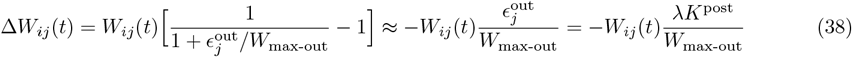

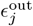 is a known quantity, given that outgoing homeostasis guarantees that before the last Hebbian update 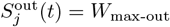, and given that at every timestep there are *K*^post^ active neurons, so the total amount of extra outgoing connectivity in *j* is *λK*^post^ upon neuron *j* firing.

Therefore, the mean update in synapse *ij* depends on the probability of pre-post coincidence (Hebbian update, Eq. (36)) and the probability of presynaptic neuron *j* firing (homeostatic update, Eq. (38)):

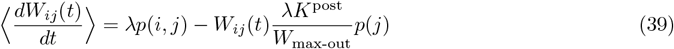

which can be interpreted as *W*_*ij*_ low-pass filtering conditional firing probabilities, (average brackets are dropped from now on, and everything represents mean field):

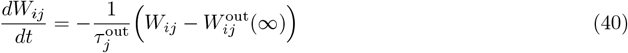

With

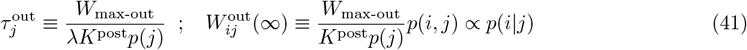

Therefore, the explicit mean synaptic trajectory of synapse *ij* can be written as:

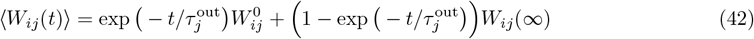

For completion, we re-write the mean-field dynamics without notation simplification:

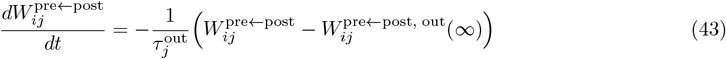

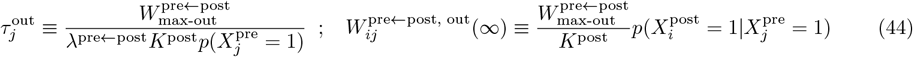

### Incoming homeostasis

Similarly, assuming on connections are initialized with 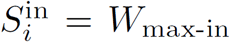 and incoming homeostasis, one can obtain:

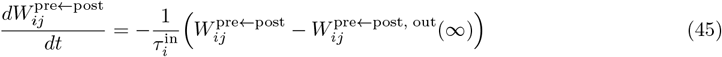

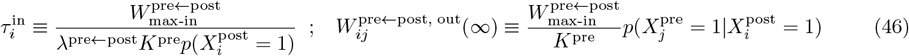

## Acknowledgments

We would like to thank Jesse Geerts, as well as the rest of the members of the Clopath Lab, for discussions and comments on earlier versions of the manuscript.

## Code Availability

All code supporting the findings of this study is available at https://github.com/albesagonzalez/homeostatic-binary-networks

